# Pairing two growth-based, high-throughput selections to fine tune conformational dynamics in oxygenase engineering

**DOI:** 10.1101/2020.05.22.111575

**Authors:** Sarah Maxel, Linyue Zhang, Edward King, Derek Aspacio, Ana Paula Acosta, Ray Luo, Han Li

**Author notes:** These authors contributed equally (Sarah Maxel, Linyue Zhang, Edward King).

## Abstract

Cyclohexanone monooxygenases (CHMO) consume molecular oxygen and NADPH to catalyze the valuable oxidation of cyclic ketones. However, CHMO usage is restricted by poor thermostability and stringent specificity for NADPH. Efforts to engineer CHMO have been limited by the sensitivity of the enzyme to perturbations in conformational dynamics and long-range interactions that cannot be predicted. We demonstrate a pair of aerobic, high-throughput growth selection platforms in *Escherichia coli* for oxygenase evolution, based on NADPH or NADH redox balance. We utilize the NADPH-dependent selection in the directed evolution of thermostable CHMO and discover the variant CHMO GV (A245G-A288V) with a 2.7-fold improvement in residual activity compared to the wild type after 40 °C incubation. Addition of a previously reported mutation resulted in A245G-A288V-T415C which has further improved thermostability at 45 °C. We apply the NADH-dependent selection to alter the cofactor specificity of CHMO to accept NADH, a less expensive cofactor than NADPH. We identified the variant CHMO DTNP (S208D-K326T-K349N-L143P) with a 21-fold cofactor specificity switch from NADPH to NADH compared to the wild type. Molecular modeling indicates that CHMO GV experiences more favorable residue packing and backbone torsions, and CHMO DTNP activity is driven by cooperative fine-tuning of cofactor contacts. Our introduced tools for oxygenase evolution enable the rapid engineering of properties critical to industrial scalability.

## INTRODUCTION

Biooxygenation can provide a viable alternative to traditional means of synthetic chemistry for the selective activation of C-H bonds. Many members of the diverse oxygenase families show great potential as catalysts, including two-component flavin hydroxylases^1,2^, Cytochrome P450s^3^, and Baeyer– Villiger monooxygenases (BVMOs)^4^. Directed evolution has been widely exploited to tailor these oxygenases for desired reactions; however, its full potential has always been limited by the relatively low throughput of downstream selection technology. To address this issue there have been advancements in designing efficient libraries with smaller theoretical sizes^5^ and ultrahigh-throughput screening methods utilizing colorimetric or fluorescence sorting using unique substrate characteristics^6^, *in vitro* microfluidic assays^7,8^, and *in vivo* cell-based platforms^9^. Although these processes have facilitated the successful directed evolution of a number of enzymes, there is a need for general and accessible methods that do not require specialized reagents, proxy readout signals, or expensive equipment.

Recently, we described the construction and application of an aerobic growth-based, high-throughput selection platform for engineering NADPH-dependent oxygenases^10^. We demonstrated that this selection platform enabled rapid remodeling of the *Pseudomonas aeruginosa* 4-hydroxybenzoate hydroxylase (PobA) active site to efficiently accept a non-native substrate 3,4-dihydroxybenzoic acid (3,4-DHBA). The selected variants appear to recognize the new substrate with synergistic hydrogen bond networks, which are difficult to discover without high-throughput searching of protein sequence space. However, the key limitation of this method is that it is not compatible with engineering NADH-dependent oxygenases. While NADPH-dependent oxygenases such as most P450s naturally function in anabolism, the NADH-dependent oxygenases function in catabolism and enable the conversion of various recalcitrant substrates such as xylene^11^ and plastics^12^. Here, we report an aerobic growth-based selection for NADH activity, which complements the NADPH-dependent selection platform. Together, these two selection systems may support the full range of engineering capabilities to develop catalytically important oxygenases.

As a proof-of-concept, we applied the two high-throughput, growth-based selection platforms to an NADPH-dependent BVMO, *Acinetobacter sp.* cyclohexanone monooxygenase (*Ac* CHMO). This class of monooxygenases is known for catalyzing the oxidation of (cyclic) ketones into esters and lactones, and have been studied for their industrial potential in the production of nylon monomers^13^, (Z)-11-(heptanoyloxy)undec-9-enoic acid^14^, and prostaglandins^15^. *Ac* CHMO has been extensively characterized in particular due to its versatile substrate scope and superior activity. However, fundamental limitations to large-scale use of this enzyme are its well documented temperature instability^16,17^ and poor activity with NADH. NADH is the preferred cofactor for bioproduction processes when considering cost, stability, and regeneration systems^18^; however, BVMOs are notoriously averse to cofactor specificity switching with existing methods^5,18,19^.

Through a single round of NADPH-dependent selection at an elevated temperature, we obtained a variant, CHMO GV (A245G-A288V), from a large library generated by error-prone PCR. GV displayed superior thermostability after a 40 °C incubation by retaining a 2.7-fold residual activity when compared to the wild type (CHMO WT). Through a single round of NADH-dependent selection, we obtained a variant, CHMO DTNP (S208D-K326T-K349N-L143P), with a roughly 21-fold improvement of catalytic efficiency *(k_cat_/K_M_*) with NADH over NADPH in comparison to CHMO WT. Importantly, molecular dynamics (MD) simulation suggest that the selected mutations may function by tuning the conformational dynamics of the protein and the cofactors, which could not be readily predicted in structure-guided protein design. For example, the key mutation L143P in CHMO DTNP emerged as a spontaneous mutation outside the three rationally picked positions (S208, K326, K349) for site-saturated mutagenesis. Although L143P does not directly interact with NADH, MD analysis suggests that it tunes the conformation of flavin adenine dinucleotide (FAD) in CHMO DTNP, allowing more efficient hydride transfer from NADH. This effect would not be evident through analysis of static models, highlighting the difficulty of engineering functionally innovative variants through rational design and the key role of MD in examining protein conformational landscapes.

Oxygenases are promising catalysts, but difficult to engineer due to their highly complex behavior in sampling multiple conformational states during catalysis^20–23^. We envision that the tools developed here will make the conformational dynamics a more tunable element in oxygenase engineering.

## RESULTS AND DISCUSSION

### Development of NADH-dependent selection platform

Both the NADPH- and NADH-dependent selection platforms utilize the principle of cofactor redox balance, where NAD(P)H accumulates in engineered *E. coli* cells to a toxic level. The cell growth is restored by expressing a heterologous oxygenase mutant with the desired activity^10,24^ to alleviate imbalance (Figure 1). As previously reported, aerobic NADPH accumulation in strain MX203 (Table S1) was achieved by channeling glucose metabolism through the pentose phosphate pathway and eliminating native NADPH sinks^10^ (Figure 1A). However, accumulating NADH in aerobic conditions represents a different plethora of challenges, since numerous metabolic pathways funnel NADH through the respiration chain. These challenges were tackled here by disrupting multiple respiratory chain components: *ndh* (encoding NADH:quinone oxidoreductase II), *nuoF* (encoding subunit of NADH:ubiquinone oxidoreductase I), and *ubiC* (encoding chorismate lyase, which catalyzes the first committed step in the biosynthesis of ubiquinone, an electron carrier in the respiration chain). In addition, fermentative pathways were blocked through disrupting *adhE*, *ldhA*, and *frdBC*^25^, and conversion of NADH to NADPH was diminished through disrupting *pntAB* (Figure 1B). Notably, a viable strain containing the above-mentioned knockouts could not be made without simultaneously expressing a water-producing NADH-oxidase from *Lactobacillus brevis* (*Lb* Nox)^26^, indicating the successful establishment of high NADH stress to a growth-inhibitory level. *Lb* Nox was expressed under an arabinose-inducible promoter (on plasmid pLS101, Table S1), so that its expression can be turned off during selection of desired oxygenases. The final NADH accumulation strain harboring a tightly controllable *Lb Nox* was named MX304 (Table S1).

**Figure 1.**
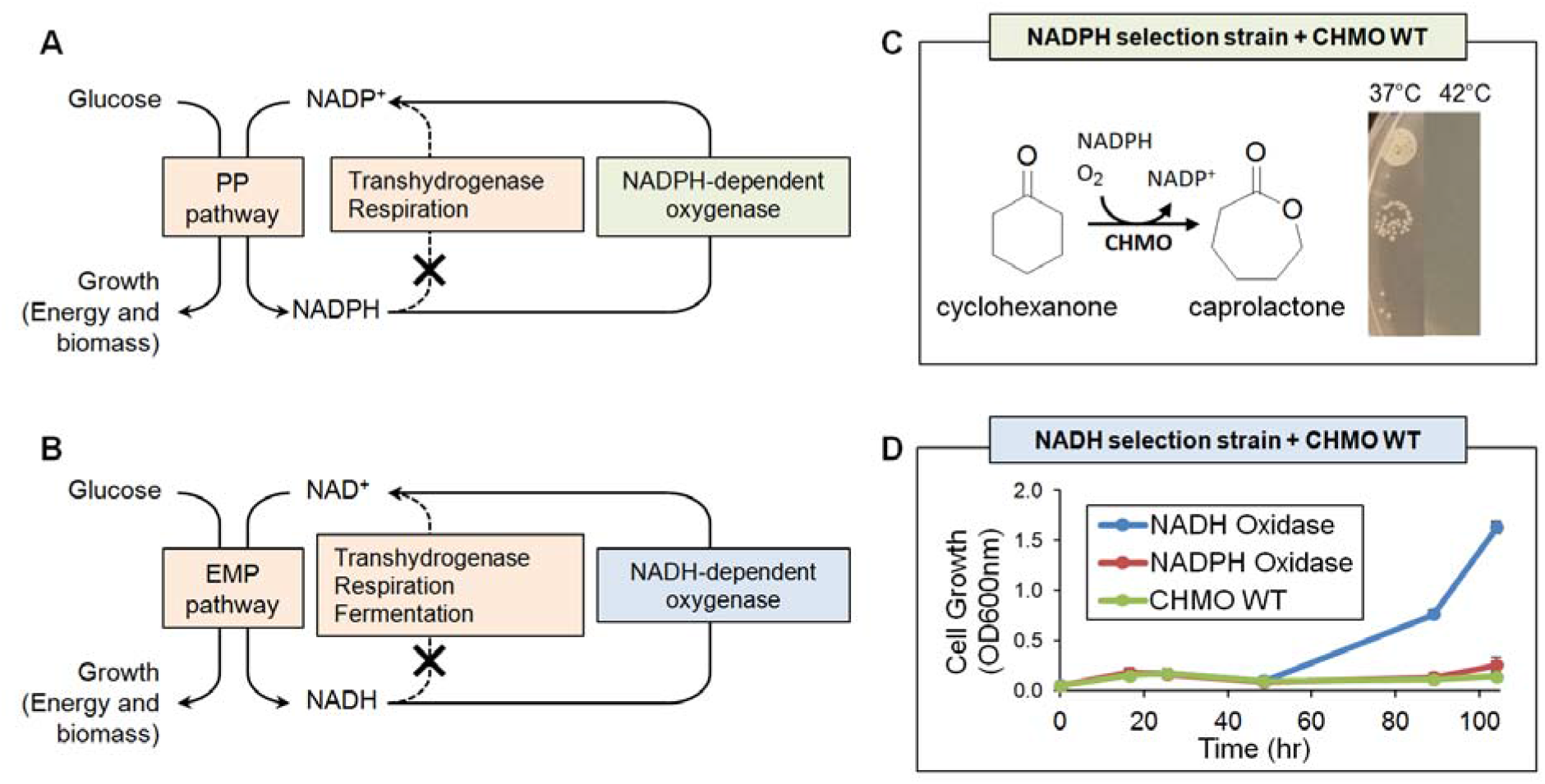
Redox balance-based selection platforms for NADPH- and NADH-dependent oxygenases. **A)** and **B)** A heterologous oxygenase forms a closed redox loop with available glucose-metabolizing pathways, pentose phosphate (PP) pathway and Embden–Meyerhof–Parnas (EMP) pathway, respectively, to enable cell growth. Natural pathways to recycle NAD(P)H are disrupted. **C)** Wild type *Ac* CHMO (CHMO WT) consumes NADPH while converting cyclohexanone to caprolactone, and restored growth of the NADPH selection stain (MX203) at 37 °C but not at 42 °C, suggesting its poor thermostability. **D)** Growth of NADH selection strain (MX304) is rescued by an NADH oxidase, but not by an NADPH oxidase or CHMO WT. Values are an average of at least three replicates, and the error bars represent one standard deviation.

We tested the growth response of these two selection strains with wild type *Ac* CHMO (CHMO WT, on plasmid pLS201), which is NADPH-dependent. As expected, CHMO WT can restore growth of the NADPH-dependent selection strain (MX203) at 37 °C in minimal medium on glucose, but failed to restore the growth of the NADH-dependent selection strain (MX304, with its arabinose-inducible copy of *Lb Nox* turned off) in the same conditions (Figure 1C, D). Importantly, no growth was observed at 42 °C for MX203 with CHMO WT (Figure 1C), suggesting that the activity level of CHMO WT is significantly reduced at elevated temperatures, which is consistent with its well documented poor thermostablity^16,17^.

We used *Lb Nox* and its NADPH-dependent counterpart, *TP Nox*^10,27^ introduced on plasmids pLS206 and pLS207, respectively (Table S1), to further validate the newly developed NADH-dependent selection platform. As expected, only *Lb* Nox, but not *TP* Nox rescued the growth of MX304 (with its arabinose-inducible copy of *Lb Nox* turned off) in both liquid and solid media (Figure 1D, Figure S1). These results confirm that the selection platform exhibits the desired cofactor specificity and can be used with flexible experimental setup.

### Identification of CHMO variants with increased thermostability

To address poor thermostability, two main strategies have been approached. First, there have been extensive efforts to expand the substrate scope of BVMOs which have good stability but relatively narrow substrate ranges^15,28–30^. Second, protein engineering efforts have been applied to make the broad substrate-range *Ac* CHMO less labile, which include introducing computationally predicted disulfide bonds^17,23^ and creating chimeric enzymes between *Ac* CHMO and its more thermostable homologs^31^. Here, we sought to leverage the high throughput of growth-based selection and mine beneficial mutations from a larger protein sequence space generated by error-prone PCR. The rationale is that stabilizing mutations can be found throughout the protein structure in regions associated with unfolding steps^32^; however, these regions can be challenging to predict.

A mutation rate of one to two mutations per variant was targeted during the construction of CHMO library (pLS203. Table S1)^33^. A library size of roughly 2.4 × 10^7^ variants was subjected to selection in strain MX203 on agar plates with 2 g/L D-glucose in M9 minimal medium and 2 g/L cyclohexanone at 42 °C. The selection resulted in a single colony with superior growth, while ~18 additional colonies were observed with significantly slower growth. Due to the drastic growth difference between colonies, only the single fast growing colony was extensively characterized. Sequencing of this variant revealed two mutations, A245G and A288V, previously unidentified as relevant to protein stability. The double mutant *Ac* CHMO A245G-A288V (CHMO GV) demonstrated reliable growth restoration of strain MX203 at 42 °C (Figure 2A) and retained superior residual activity after prolonged exposure to a range of elevated temperatures when evaluated alongside CHMO WT *in vitro* (Figure 2B). After a 10-minute incubation at 40 °C, WT retained only 19% activity while GV retained 52%. In industry, thermotolerance between 30-40 °C is preferred. After incubation in this range, CHMO GV is roughly 1.2-4.4 fold more active than CHMO WT. While having improved thermostability, CHMO GV also retains comparable catalytic efficiency compared to CHMO WT (Table S2).

**Figure 2.**
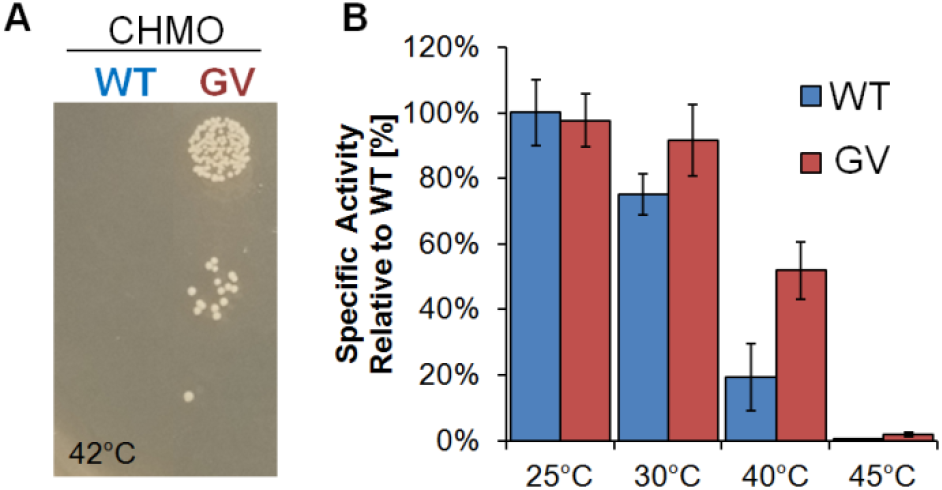
Analysis of CHMO GV with improved thermostability. **A)** At 42 °C, CHMO GV, but not WT, restores growth of the selection strain MX203. **B)** After 10 minutes of high temperature exposure, thermostable CHMO GV retains higher activity compared to WT. Values are an average of two or three replicates, and the error bars represent one standard deviation.

After incubation at 45 °C, CHMO WT is completely deactivated while CHMO GV is still 2% active. A CHMO variant which is more stable at 45 °C was obtained by adding T415C, a previously identified stability enhancing mutation^34^ (Figure S2). This suggests that improvements can be continually accessed by iterative rounds of random point mutation. While the current platform constructed in mesophilic *E. coli* is limited by growth inhibition at temperatures exceeding 42 °C, construction of similar selection strains in a heat resistant *E. coli* strain with a maximum growth rate at 48.5 °C ^35^ or thermophilic bacteria such as *Geobacillus thermoglucosidasius* may allow facile identification of increasingly thermostable variants.

### The effect of thermostability enhancing mutations on protein dynamics

Since there are no crystal structures of *Ac* CHMO with cofactors bound available, we constructed a homology model for the NADPH and FAD bound *Ac* CHMO structure using Rosetta CM, based on *Rhodococcus sp.* CHMO^23^, which shares 57.8% sequence identity. We ran 400 ns MD trajectories of CHMO GV and CHMO WT, and featurized the trajectories on minimum heavy atom distance from residues within 5 Å of position 245 or 288, followed by dimensionality reduction with principle component analysis (PCA). The conformations are discretized with K-means clustered in PCA space to identify the most populated metastable states for analysis (Figure S3). We visualize the frame with minimum distance to the cluster center as the representative conformations involving positions 245 and 288 (Figure 3A, B, D, E). We further extract 200 frames from the most populated cluster of each sample, and evaluate the local structure stability as a function of the local Rosetta energy, which is the sum of residue energies measuring the favorability of steric, geometric, and electrostatic interactions for all amino acids within 5 Å of positions 245 and 288 (Figure 3C).

**Figure 3.**
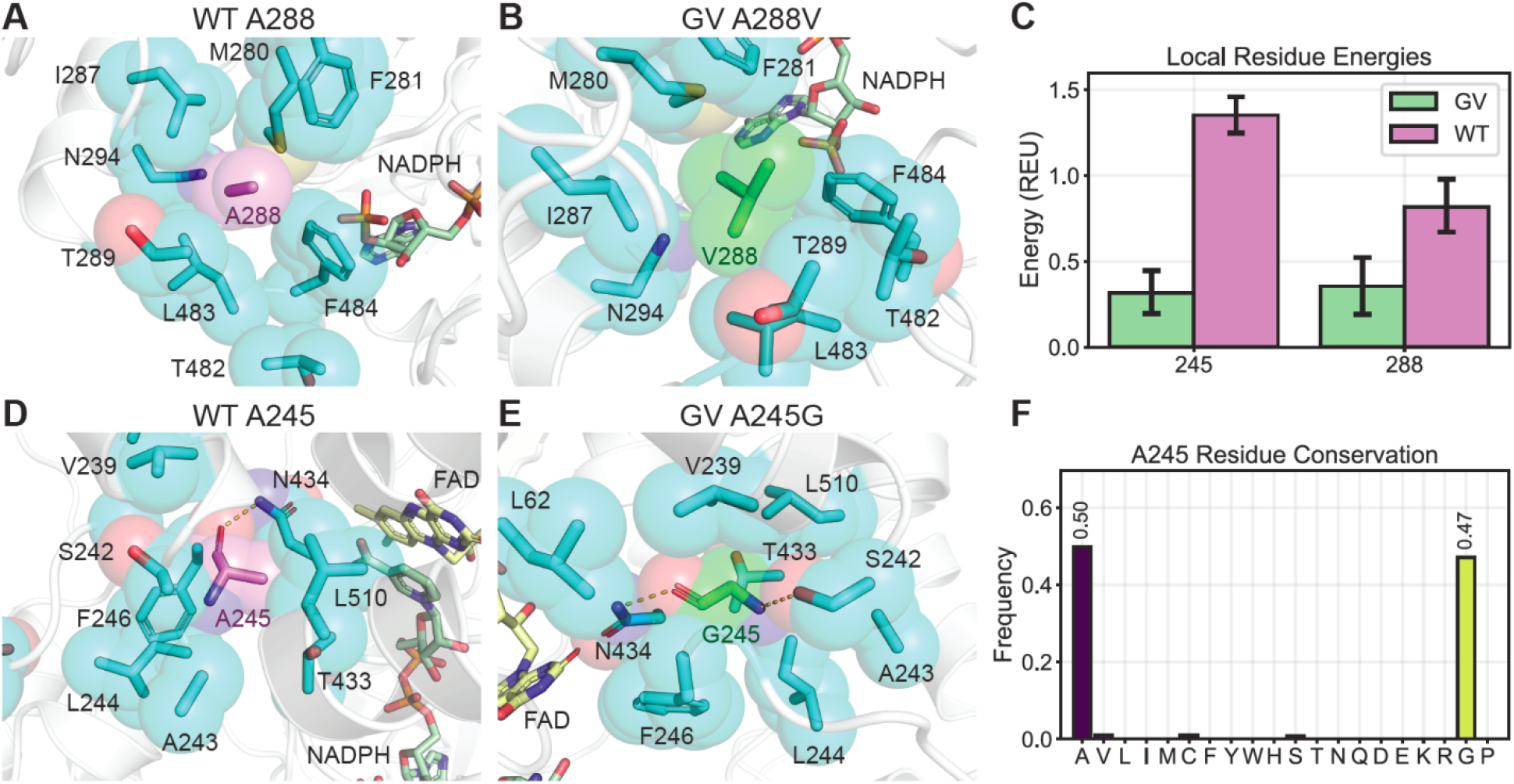
Molecular dynamics (MD) conformational analysis indicates that more optimal residue packing and backbone torsion underlies the increased thermostability of CHMO GV. **A)** and **B)** At position 288, valine has greater volume and surface area to maximize hydrophobic contact with surrounding residues. **C)** Rosetta energy evaluation of snapshots from the MD trajectories indicates that interacting residues surrounding the mutated positions experience more favorable contacts in CHMO GV compared to WT. **D)** and **E)** At 245 position, glycine is better enveloped by nearby residues, and has greater backbone torsion to form an additional backbone hydrogen bond. **F)** Measurement of residue frequencies from sequence alignment of *Ac* CHMO homologs shows that A and G are highly conserved at position 245. Values are an average of 200 frames and error bars represent 95% confidence intervals with 1,000 bootstrap iterations.

In CHMO WT, A288 leaves excess space for water molecules to enter and causes undesired sidechain flexibility (Figure 3A). In contrast, A288V in CHMO GV completely fills the small void created by the residues nearby and provides a greater nonpolar surface area to encourage closer packing (Figure 3B). As a result, CHMO GV exhibits greater stability at position 288, scoring 0.35 ± 0.2 Rosetta Energy Units (REU), compared to 0.81 ± 0.2 REU for CHMO WT (Figure 3C). At 245 position, the reduced bulk from A245G allows the surrounding residues to pack more tightly in CHMO GV with fewer gaps for solvent to enter (Figure 3D, E). More importantly, the increased torsion flexibility from glycine permits A245G in CHMO GV to contort and form backbone hydrogen bonds with N434 and S242. The more restricted torsions prevent A245 in CHMO WT from satisfying both backbone hydrogen bonds simultaneously (Figure 3D, E). These interactions are reflected by the local energy scores (Figure 3C), with CHMO GV scoring more favorably than CHMO WT at position 245 (0.32 ± 0.1 REU versus 1.35 ± 0.1 REU).

Interestingly, bioinformatic analysis of natural residue diversity at A245 in *Ac* CHMO homologs indicates that alanine and glycine are sampled in 97% of homologous sequences, with glycine in 47% of sequences and alanine in 50% (Figure 3F). Our selection method which had access to the full set of amino acids resolved to the same highly conserved residue collection as Nature, indicating that there is a strong selective pressure to maintain small amino acids at position 245, and that our selection explores sequence space in a manner compatible with natural evolution.

### Directed evolution of CHMO for NADH-dependent activity

BVMOs cycle between the “open” and “closed” conformations during catalysis, with the cofactor NADP(H) not only participating in the reaction but also playing an important role in coordinating the conformational change^23^. Therefore, we hypothesized that to switch the cofactor specificity of *Ac* CHMO, it would be crucial to target sites that guide the conformational changes accompanying catalytic cycles, in addition to the ones that directly discriminate the 2’-phosphate of NADPH. Specifically, we chose four positions, K326, K349, S208, and T378. These sites were also subjected to extensive rational design efforts by Beier *et al.* to improve NADH-dependent activity^18^. However, only a fraction of the possible sequence space (20^4^ = 1.5 × 10^5^) was searched by rational engineering strategies.

We incorporated NNK degenerate codons at the four sites mentioned above to yield a CHMO library (pLS209, Table S1). The library size constructed was estimated to be 2.5 × 10^7^ independent transformants. Once the diversity of the library was confirmed by sequencing, pLS209 was transformed into the selection strain MX304 and the selection was conducted on M9 selection agar plates containing 2 g/L D-glucose and 2 g/L cyclohexanone at 30 °C for up to 120 h. Interestingly, many colonies harbored the same variant, CHMO DTNP (S208D-K326T-K349N-L143P). This variant retained the wild type residue at position T378 that is typically conserved across the BVMO family^18^, and contained an unanticipated mutation L143P which possibly originates from spontaneous error in PCR or *in vivo* plasmid replication.

We characterized the kinetic parameters of the CHMO variants towards different cofactors (Table 1). Compared to wild type, CHMO DTNP exhibited ~2-fold increase in catalytic efficiency *(k_cat_/K_M_*) for NADH and ~ 10-fold decrease for NADPH, which collectively give a 21-fold cofactor specificity switch. Individual characterization of *k_cat_* and *K_M_* was not achieved because the enzymes could not be saturated in the range of cofactor concentrations tested. The improvement we achieved on NADH-dependent activity is substantial, given BVMOs well-documented recalcitrance to efforts aiming to increase NADH activity^19^. To benchmark CHMO DTNP, we compared it to the CHMO variant PEHR (S186P-S208E-K326H-K349R), which was reported to have the largest cofactor specificity switch and the best performance in NADH-dependent biotransformation among the rationally engineered variants^18^. The kinetic parameter analysis shows that our selected variant DTNP has significantly improved catalytic efficiency for NADH (45.8 ± 5.51 mM^-1^s^-1^ for DTNP versus 21.5 ± 1.69 mM^-1^s^-1^ for PEHR. Tables 1, S3).

**Table 1.** Kinetic characterization of CHMO variants

| CHMO variants | $k_{cat}/K_m$ (mM<sup>-1</sup>s<sup>-1</sup>) | |
| --- | --- | --- |
|  | NADH | NADPH |
| WT | $20.8 \pm 3.51$ | $400 \pm 25.8$ |
| DTNP (S208D-K326T-K349N-L143P) | $45.8 \pm 5.51$ | $41.5 \pm 6.04$ |
| DTN (S208D-K326T-K349N) | $26.8 \pm 0.38$ | $47.3 \pm 2.02$ |

Furthermore, the kinetic characterization shows that the spontaneous mutation L143P is critical to increased NADH-dependent activity (Table 1), which is consistent with the fact that CHMO DTN (S208D-K326T-K349N) was never discovered in growth selection. Contrastingly, the catalytic efficiency for NADPH is minimally impacted by this spontaneous mutation (Table 1, comparing CHMO DTNP and DTN), suggesting that L143P is specifically involved in NADH-dependent catalysis.

### The effect of cofactor specificity-switching mutations on protein and cofactor dynamics

Rosetta modeling suggests that S208D and K349N function cooperatively to recognize NADH, with K349N supporting the S208D loop through backbone hydrogen bond and S208D forming a novel bidentate hydrogen bond to the NADH adenosine ribose that resembles interactions seen in native NADH specific proteins (Figure 4A, B). The contribution of K326T and L143P are not evident from static structural analysis since they do not directly contact NADH (Figure 4A, C, D), which motivated further analysis through MD simulations.

**Figure 4.**
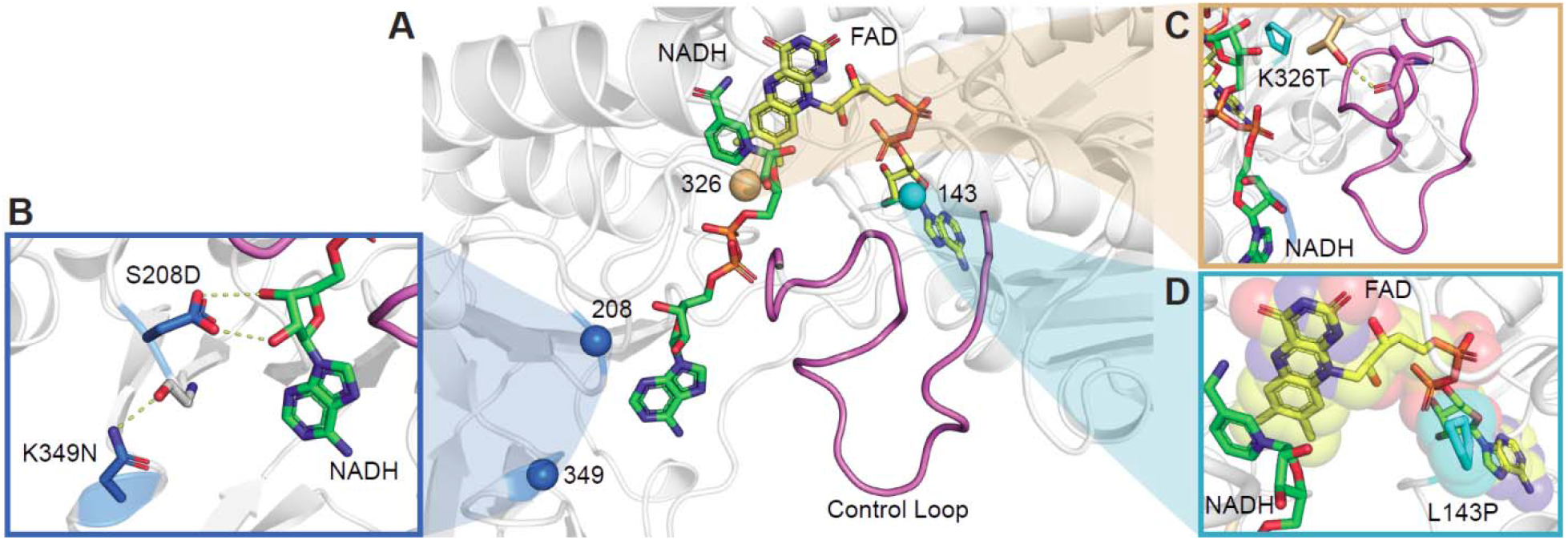
Homology Model of CHMO DTNP. **A)** Overview of the cofactor binding pocket, NADH (in green) is pressed on one end by the control loop (purple). FAD (yellow) is held opposite of the NADH, and positions of the selected mutations are represented as spheres. **B)** K349N supports the loop that S208D sits on, and S208D makes a bidentate hydrogen bond to the NADH adenosine ribose. **C)** K326T initiates a hydrogen bond to the control loop at the backbone carbonyl of A487. The additional hydrogen bond may cause the control loop to favor maintaining the catalytically relevant, closed conformation. **D)** L143P impacts the conformations that FAD is able to adopt.

K326T contacts the “control loop” through A487 (Figure 4A, C). The control loop is observed to be ordered in some CHMO crystal structures but disordered in others^21,23,36^, suggesting that its flexibility varies depending on the enzyme’s position in catalytic cycle. Specifically, it is hypothesized that the control loop must become rigid to hold the NADPH cofactor and cyclohexanone during the hydride transfer stages^21^. We analyzed the flexibility of the control loop through α-carbon root mean square fluctuation (RMSF), and plot the difference between CHMO DTNP (bound with NADH) and CHMO WT (bound with NADH or NADPH, respectively) (Figure 5). The results show that CHMO DTNP with NADH bound maintains greater rigidity over the control loop in comparison to the WT with NADH bound and the level of control loop rigidity in CHMO DTNP with NADH bound is nearly identical to that of the CHMO WT binding NADPH. These results support the role of K326T in stabilizing the control loop which clamps on the otherwise loosely bound cofactor NADH.

**Figure 5.**
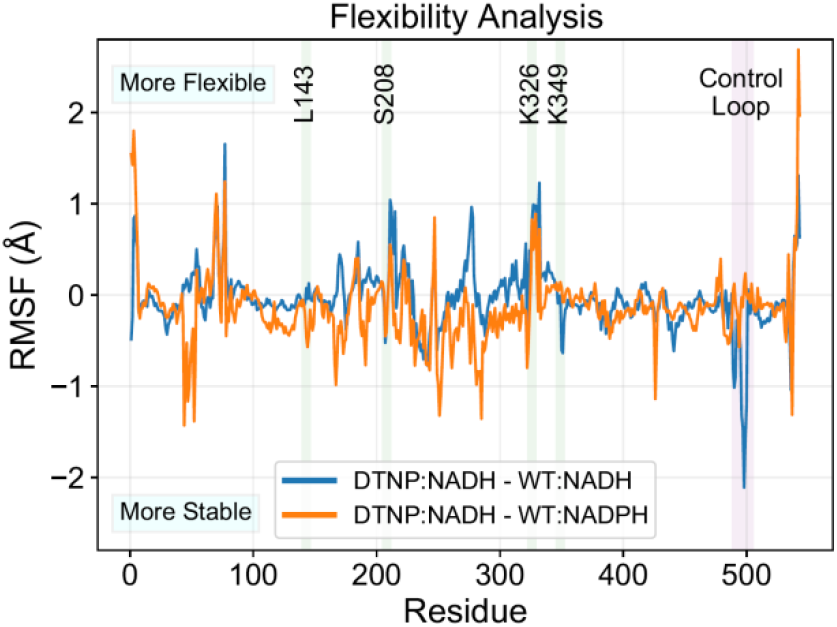
Root mean square fluctuation (RMSF) analysis of CHMO DTNP. Results are plotted as the difference in RMSF between DTNP (with NADH bound) and WT (with NADH or NADPH bound, respectively). The control loop is found to have greater stability with DTNP:NADH, comparable to the native condition of WT:NADPH. The inactive pairing of WT:NADH exhibits high flexibility at the control loop.

L143P contacts FAD (Figure 4D, 6). Therefore, we hypothesized that this mutation is involved in positioning FAD and in turn affects hydride transfer from the nicotinamide cofactors to FAD. We first compared the hydride transfer distances (the distance between nicotinamide C4 and FAD N5) when CHMO WT utilizes different cofactors, and confirmed that higher activity is linked to a shorter hydride transfer distance (4.6 ± 0.5 Å for NADPH versus 6.5 ± 1.0 Å for NADH, Figure 6A)^23^. Interestingly, the hydride transfer distance when using NADH as the cofactor was much shorter in CHMO DTNP compared to in CHMO DTN (Figure 6B, C), which is consistent with our hypothesis that L143P facilitates more efficient hydride transfer from NADH.

**Figure 6.**
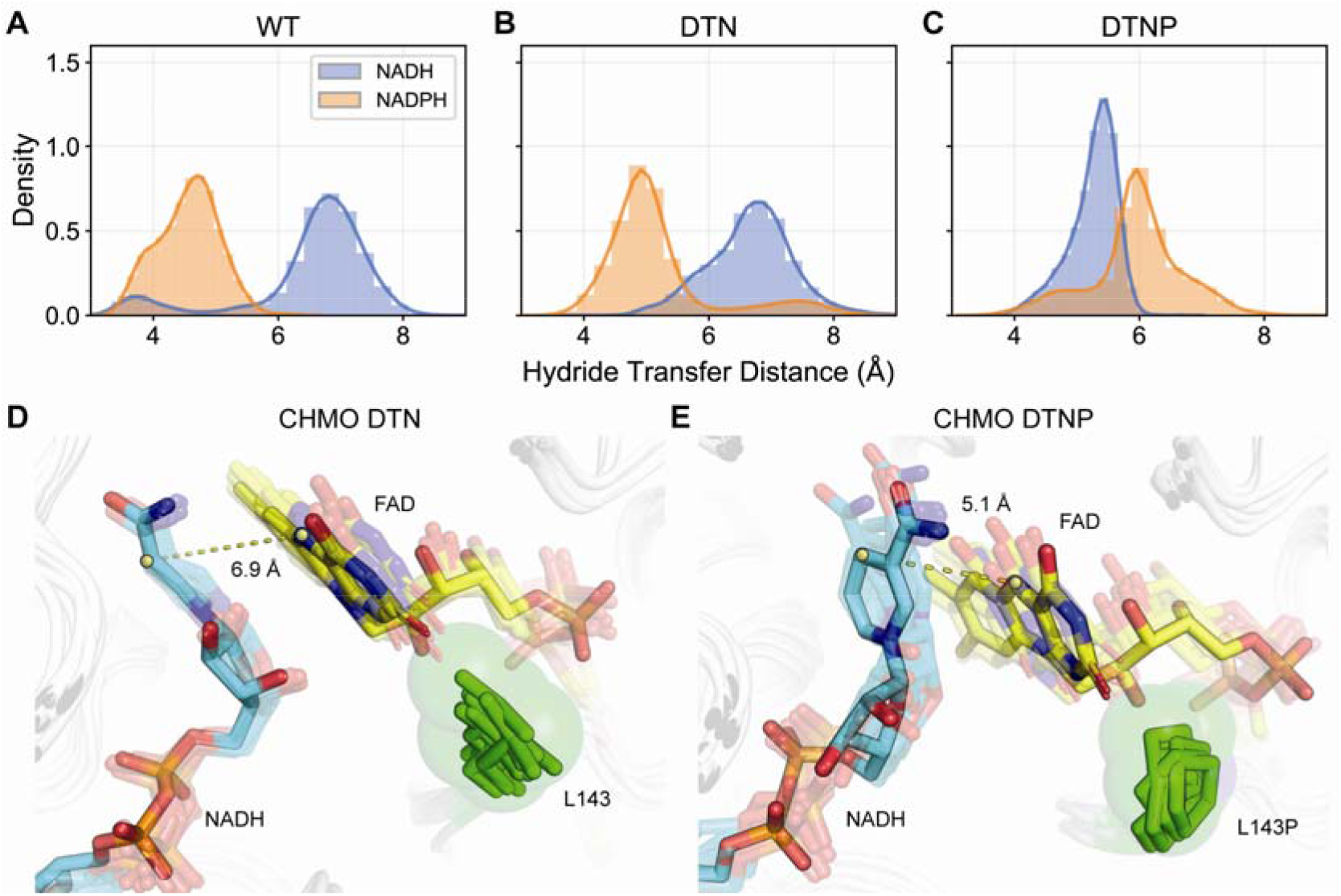
Evaluation of hydride transfer efficiency in CHMO variants. Shorter distances between NAD(P)H C4 and FAD N5 contributes to enhanced catalysis. **A, B)** WT and DTN are both marked by NADPH sampling distances ~5 Å, while the less active NADH samples distances >6 Å and is too remote to engage in hydride transfer. **C)** DTNP displays the opposite arrangement, with NADH sampling closer distances than NADPH. **D)** L143 in DTN firmly packs under the flavin ring, blocking the flavin from moving closer to nicotinamide, resulting in suboptimal hydride transfer distance. **E)** L143P presses against the FAD ribitol rather than contacting the flavin. This anchors the FAD core and allows the flavin head to rotate in response to changes in the nicotinamide positioning to sustain closer contact. The dashed lines show representative distances between the nicotinamide C4 and FAD N5.

To understand the role of L143P in influencing hydride transfer, we identified metastable FAD conformations in CHMO DTN (with NADH bound) and CHMO DTNP (with NADH bound) based on minimum heavy atom distance to residues within 4 Å and performed PCA with K-means clustering (Figure S4). Conformations from the most populated cluster of each sample were compared and indicate that the native leucine packs underneath the flavin (Figure 6D), limiting flavin’s flexibility to move closer to the nicotinamide. This limited flexibility is likely advantageous under the native condition of NADPH binding to CHMO WT, resulting in precise organization of the flavin for hydride transfer. However, upon mutation at S208D and K326T which contact the adenosine end of the cofactor, the binding pose of the cofactor is altered, which requires the flavin to adjust its position accordingly. The L143P proline in CHMO DTNP appears to pack along the FAD ribitol instead of directly against flavin (Figure 6E). This may allow twisting motion of FAD to orient the flavin toward the NADH nicotinamide ring.

## CONCLUSION

In summary, we established a growth-based, high-throughput selection platform in *E. coli* to facilitate the directed evolution of NADH-dependent oxygenases, which complements the NADPH-dependent systems we developed previously^10^. We used these two selections in parallel to engineer a monooxygenase, CHMO, toward a more viable catalyst in large-scale processes. Specifically, we sought to address fundamental limitations characteristic to this biocatalyst: poor thermostability and strict cofactor preference for NADPH, which is more expensive than NADH. Despite engineering efforts to tackle both limitations previously^18,31,34,37^, we hypothesized that exploration of a wider sequence space would enable the discovery of innovative variants. First, selection of an error prone PCR library at an elevated temperature enabled the rapid isolation of thermostable CHMO GV which is 1.2 and 4.4-fold more active than CHMO WT after exposure to industrially relevant temperatures, 30 and 40 °C respectively. Second, selection of a four-residue, site-saturated mutagenesis library obtained CHMO DTNP with 21-fold cofactor specificity switch from NADPH to NADH, and significantly better catalytic efficiency for NADH compared to previously reported variant made by rational design^18^. Interestingly, the contributions of the selected mutations are not immediately evident from inspection of static structure. For example, CHMO GV contains a mutation to glycine which is typically associated with reduced thermostability due to added flexibility^38^; CHMO DTNP harbors a spontaneous mutation L143P which falls outside the targeted sites deemed important in cofactor recognition. MD simulations highlight that most of the mutations appear play a role in fine tuning the conformational dynamics of enzyme and cofactor, which are subtle effects that are challenging to design without a high-throughput method. Conformational dynamics is particularly critical for catalysis in oxygenases^20–23^. We envision the wide application of our selections in directed evolution of these complex and powerful enzymes.

## SUPPORTING INFORMATION

Experimental and computational methods, plasmids and strains used in this study (Table S1), detailed kinetic analysis of CHMO variants (Table S2), growth phenotypes of *E. coli* strains with modified NADH metabolism (Figure S1), comparison of residual specific activity of CHMO WT and GV to previously described mutation T415C and a combined variant (A245G-A288V-T415C) after 45 °C incubation (Figure S2), free energy landscapes for CHMO GV and WT (Figure S3), and free energy landscapes for CHMO DTN, DTNP, and WT with NADH or NADPH (Figure S4).

## AUTHOR INFORMATION

**Corresponding Author**

* Han Li

## AUTHOR CONTRIBUTION

S.M., L.Z., and H.L. designed the experiments, S.M., L.Z., D.A., and A.P.P performed the experiments and analyzed the results, E.K. performed Rosetta and MD modeling, E.K. and R.L. analyzed the modeling results. All authors wrote the manuscript.

## NOTES

The authors declare no competing financial interest.

## ACKNOWLEDGMENT

H.L. acknowledges support from University of California, Irvine, the National Science Foundation (NSF) (award no. 1847705), and the National Institutes of Health (NIH) (award no. DP2 GM137427). S.M. acknowledges support from the NSF Graduate Research Fellowship Program (grant no. DGE-1839285). D.A. acknowledges support from the Federal Work Study Program funded by the U.S. Department of Education.

## Notes

### Competing Interest Statement

The authors have declared no competing interest.

