## Supporting information for "Pairing two growth-based, high-throughput selections to fine tune conformational dynamics in oxygenase engineering"

Departments of Chemical and Biomolecular Engineering[†], Molecular Biology and Biochemistry[§], Biomedical Engineering[|], University of California, Irvine. [‡]These authors contributed equally.
*To whom correspondence should be addressed

**Supporting Information**

**A. Methods**

**B. Supplemental Tables and Figures**

Table S1 Strains and Plasmids used in this study.

Table S2 Kinetic parameters of CHMO WT and GV.

Table S3 Kinetic parameters of CHMO PEHR.

Figure S1 Modified metabolisms of *E. coli* strains displayed distinct growth phenotypes in M9 Selection Media with glucose as carbon source.

Figure S2 Residual specific activity of CHMO GVC (A245G-A288V-T415C) after ten-minute incubation at 45 °C.

Figure S3 Free energy landscapes for CHMO GV and WT.

Figure S4 Free energy landscapes for CHMO DTN, DTNP, and WT with NADH or NADPH bound.

**C. References**

### A. Methods

**Media and Growth Conditions**. Cloning was carried out with *E. coli* XL-1 blue and protein expression was performed with *E. coli* BL21 (DE3) or MX203 as indicated. All *E. coli* were cultured in 2xYT containing 16 g/L Tryptone, 10 g/L Yeast Extract and 5 g/L NaCl unless otherwise noted. M9 Wash Buffer consisted of 1 mM $MgSO_4$, 0.1 mM $CaCl_2$, trace metal mix A5 with Co ($H_3BO_3$ 2860 μg/L, $MnCl_2 \cdot 4H_2O$ 1810 μg/L, $ZnSO_4$ $7H_2O$ 222 μg/L, $Na_2MoO_4$, $2H_2O$ 390 μg/L, $CuSO_4$, $5H_2O$ 79 μg/L, $Co(NO_3)_2$ $6H_2O$ (49 μg/L), and BD Difco M9 salts ($Na_2HPO_4$ 6.78 g/L, $KH_2PO_4$ 3g/L, NaCl 0.5 g/L, $NH_4Cl$ 1 g/L). M9 Selection Media shared the same composition of M9 Wash Buffer with the inclusion of 2 g/L D-glucose, 0.01 g/L thiamine, 0.04 g/L $FeSO_4 \cdot 7H_2O$. For solid media M9 Selection Plates, 15 g/L agar was added in addition to M9 Selection Media composition. In overnight cultures substrate cyclohexanone was added at a concentration of 1 g/L and in growth restoration experiments 2 g/L was added from a concentrated stock when appropriate. Concentrations for antibiotic selection were 100 mg/L for ampicillin, 50 mg/L for kanamycin, 50 mg/L for spectinomycin, and 10 mg/L for tetracycline. All strains were cultured at 37 ℃ with 250 rpm agitation unless otherwise noted. Induction in growth experiments and overnights was initiated with final concentrations of 0.01% arabinose for strains with $P_{BAD}$ promoter and 0.05 mM IPTG for strains with $P_{lac}$ promoter. Non-selection media for NADH strains MX302, MX303, and MX304 was supplemented with 0.1% arabinose and 2 g/L acetoin to provide NADH consuming outlets by heterologous NADH oxidase and endogenous gldA activity[1], respectively.

**Strain Construction.** Construction of strain MX203 was described previously[2]. *E. coli* strain JCL166 served as starting strain for NADH strain MX304 construction[3]. Plasmid pCP20 was used to eliminate kanamycin resistance in between knockouts performed[4]. Knockouts Δ*nuoF*, Δ*ndh*, Δ*ubiC*, and Δ*pntB* on JCL166 were generated using the P1 phage transduction method[5]. Keio collection strains[4] JW229-3 (*ΔnuoF::kan*), JW1095-1 (*Δndh::kan*), JW5713-1 (*ΔubiC::kan*), and JW1594-1 (*ΔpntB::kan*) served as donors for the generation of P1 lysate with gene knockout cassettes containing a kanamycin resistance marker. Construction of strain MX301 (JCL166 Δ*nuoF* Δ*ndh::kan*) resulted in a strain that demonstrated poor growth in even rich media. Outside of expressing heterologous genes to restore growth, strains will evolve under stressful conditions; therefore, suitable growth conditions were necessary to maintain the strain in non-selection conditions. Strictly regulated NADH consumption strategies were achieved via expression of *Lb* NOX on a tightly controlled arabinose inducible plasmid (pLS101). Supplementation of pLS101 and 0.1% arabinose was maintained in all further strain variants to provide NADH sink in non-selection culturing to preserve growth phenotype of the strains. All knockouts were confirmed by MyTaq Colony PCR (Bioline).

**Plasmid Construction.** All PCR fragments were generated using PrimeSTAR Max DNA Polymerase (TaKaRa) unless otherwise noted. Splicing-by-overlap extension (SOE) PCR and degenerate codon PCR was conducted using KOD Xtreme Hot Start DNA Polymerase (Novagen). The *Lactobacillus brevis nox* gene was amplified from genomic DNA from *Lactobacillus brevis* strain 118-8 [ATCC® 367™]. After PCR and gel extraction, the *Lb nox* gene fragment was inserted into the vector backbone (pRSF ori, Spec$^r$) using Gibson isothermal DNA assembly method[6], resulting in plasmid pLS101. Plasmid pLS102 carrying the *TP nox* gene was generated using plasmid pLS101 as a template and using site directed mutagenesis to introduce the following mutations G159A-D177A-A178R-M179S-P184R.

The *Acinetobacter sp. NCIMB 9871 chnB* gene was amplified from an *E. coli* codon optimized gBlock (IDT). After PCR and gel extraction, the *Ac chnB* gene fragment was inserted into the pRSF vector backbone which contains a 6×His tag at the C-terminus using Gibson isothermal DNA assembly method, resulting in plasmid pLS201. Plasmid pLS202 was generated using pLS201 as a template and applying site directed mutagenesis to introduce mutation T415C.

Plasmids pLS206, pLS207, and pLS208 were constructed using pLS101, pLS102, and pLS201, respectively, as templates for gene fragment amplification which were assembled with pQElac vectors. Plasmid pLS205 was generated by the addition of mutation T415C using pLS204 as a template. Plasmid pLS211 was generated by the reverting the L143P mutation to the wildtype residue using pLS210 as a template. Plasmid encoding PEHR (pLS212) was generated by addition of mutations S186P-S208E-K326H-K326R using pLS208 as a template.

**Generation of Error Prone PCR Library.** The *Ac* CHMO error prone PCR Library was cloned using the GeneMorph II Random Mutagenesis Kit (Agilent) to generate 1 to 2 mutations on gene *Ac chnB.* Using 180 ng of template DNA (provided by plasmid pLS201), the gene insert was amplified via error prone PCR with 25 reaction cycles according to manufacturer's instructions. A panel of starting template material (150 ng to 250 ng) was initially evaluated to tune final range of mutations observed. After PCR, gene fragment was digested overnight with restriction enzymes BamH-HF and SalI (NEB) in parallel with backbone fragment (NADH oxidase plasmid pLS101 provided pRSF backbone template for library construction). Library inserts and backbone fragments were ligated overnight and then transformed into ElectroMAX DH10β competent cells (Invitrogen) by electroporation. Transformed cells were rescued in 600 µL SOC medium by shaking for 1 h at 37 ℃. After rescue, cells were added to 20 mL 2xYT medium in a 250 mL baffled shake flask and 2 µL, 20 µL, and 200 µL was plated on 2xYT agar plates with appropriate antibiotics. Colonies formed on these plates were counted to estimate library size, $2.43\times 10^7$. The liquid culture was incubated at 37 ℃ for 10 h before plasmid extraction to generate the plasmid library, pLS203. Ten colonies from the library estimation plates were cultured in liquid medium for plasmid extraction and sequencing to confirm low mutation rate.

**Generation of S208-K326-K349-T378 NNK library.** The *Ac* CHMO combinatorial site-saturation mutagenesis library was cloned using primers containing NNK or MNN degenerate codons at targeted sites. PCR was used to amplify a DNA fragment from pLS208 between a forward primer beginning at position A209 and a reverse primer beginning at position A325. An additional fragment was generated by PCR of the same template using a forward primer with NNK codon at position K326 and a reverse primer with MNN codon at position K349. A final PCR fragment was amplified from the same template using primers containing a forward primer beginning at position A350 and a reverse primer containing an MNN codon at position T378. The fragments of corresponding length were purified by gel electrophoresis and three purified fragments were used as templates in an SOE PCR reaction to amplify a single fragment containing mutations at sites K326, K349, and T378. PCR was used to amplify a DNA fragment from pLS208 between a forward primer beginning at position G379 and a reverse primer containing an MNN codon at position S208. Both fragments were purified by gel electrophoresis, assembled using Gibson Isothermal Assembly, and then transformed into ElectroMAX DH10β competent cells (Invitrogen) by electroporation. Transformed cells were rescued in 600 µL SOC medium by shaking for 1 h at 37 ℃. After rescue, cells were added to 20 mL 2xYT medium in a 250 mL baffled shake flask and 2 µL, 20 µL, and 200 µL was plated on 2xYT agar plates with appropriate antibiotics. Colonies formed on these plates were counted to estimate library size. The liquid culture was incubated at 37 ℃ for 10 h before plasmid extraction to generate the plasmid library, pLS209. Ten colonies from the library estimation plates were cultured in liquid medium for plasmid extraction and sequencing to confirm diverse mutations at all four targeted sites.

**Growth Rescue Conditions.** The growth rescue condition used are as follows: Briefly, the strains tested were first cultured in 2xYT under aerobic conditions at 30 ℃ overnight with appropriate antibiotics and inducers. Next, overnight cultures were washed 3 times and re-suspended in M9 Wash Buffer. For solid growth, targeted serial dilutions of $10^6$ cells/mL, $10^5$ cells/mL, and 10 cells/mL were prepared in M9 Wash and 5 μL aliquots were dispensed in series on an agar plate of M9 Selection Media, with appropriate antibiotics and inducers. Plates were grown at designated temperatures and photos were taken to document growth progress. Plates were parafilmed to minimize evaporation of substrate and media. For liquid growth, a 0.1% (v/v) volume of washed culture was used to inoculate to an $OD_{600nm}$ of ~0.05 in 0.3 mL of M9 Selection Media, with appropriate antibiotics and inducers. Culture tubes were incubated at 30 ℃ in a rotary shaker, and $OD_{600nm}$ was measured using a 96-well plate reader.

**Transformation of libraries for selections with NADPH (MX203) or NADH strains (MX304).** To generate electro-competent *E. coli,* selection strain cells were cultured in 200 mL SOB medium with appropriate antibiotics at 30 ℃ with shaking at 250 r.p.m. until $OD_{600nm}$ reached 0.4-0.6. The culture was chilled on ice for 15 min and the cells were pelleted at 4 ℃, 4000 ×g. The cells were washed at 4 ℃ three times with 40 mL 10% glycerol solution (ice cold). After, cells were finally resuspended with 500 μL 10% glycerol solution (ice cold), and aliquoted for transformation.

The transformation was performed as follows: After electro-competent cells were prepared, 20 uL of library DNA was added to 200 μL competent cells. Cell-DNA mixture was added to four ice chilled 1 mm gap electroporation cuvettes (55 μL of per electroporation cuvette). Cells were electroporated at 2 kV, 129 Ω, 50 μF, resistance 2.5 kV; 200 uL of SOC medium was immediately added and transferred to a microcentrifuge tube at room temperature. This step was repeated twice more. Cells were rescued at 37 ℃ with shaking for 1 hour. Serial dilution of the cells was performed and then plated on 2xYT agar plates with appropriate antibiotics and addition of acetoin and arabinose as needed. After incubation at 37 ℃ overnight, colonies formed were counted to estimate transformation efficiency.

**Selection of *Ac* CHMO epPCR Library.** *E. coli* MX203 cells were transformed with the *Ac* CHMO epPCR library pLS203 via electroporation. After rescue in SOC medium for 1 hour, cultures were combined in 20 mL 2xYT with appropriate antibiotics in a 250 mL baffled shake flask. Controls were added to 5 mL 2xYT in a 50 mL conical tube (cap loose to allow increased aeration) and grown at 30 ℃ for ~5 hours or until $OD_{600nm}$ = 0.6 was reached. Subsequently, controls and the library were induced by addition of arabinose. Cultures were grown for an additional 4 hours or until $OD_{600nm}$ = ~2.2.

To prepare cells for the selection condition, 1 mL of each culture was pelleted in 2 mL microcentrifuge tubes and washed three times with M9 Wash Buffer. After wash, cells were finally re-suspended in 1 mL M9 Wash Buffer. Cells were diluted with M9 Wash Buffer to a final cell concentration of ~$10^7$ cells/mL. Twenty 100 μL aliquots of this cell suspension were plated on separate M9 Selection Plates supplemented with 2 g/L cyclohexanone and incubated at 42 ℃ for 48 hours. Expression of CHMO WT from plasmid pLS201was observed in parallel at 37 ℃ to serve as a positive control and at 42 ℃ to serve as negative control. The colony growth was monitored periodically. A single colony was observed with rapid growth comparable to positive control, while ~18 additional colonies were observed with significantly slower growth. Selected colonies were picked and re-streaked onto the fresh selection media and again incubated at 42 ℃ to obtain single colonies. Single colonies were cultured in liquid media to extract plasmids using QIAprep Spin Miniprep kit (Qiagen) to yield pLS204. Due to the drastic growth difference between colonies, only the single fast growing colony was extensively characterized.

**Selection of *Ac* CHMO NNK NADH Library.** *E. coli* MX304 cells were transformed with the *Ac* CHMO NNK NADH library pLS209 via electroporation. After rescue in SOC medium for 1 hour, cultures were combined in 20 mL 2xYT with appropriate antibiotics in a 250 mL baffled shake flask. A dilution series from culture was plated to determine transformation of individual variants: $2.47 \times 10^7$. CHMO WT (pLS208) negative control and NADH oxidase (pLS206) positive control were added to 5 mL 2xYT in a 50 mL conical tube (cap loose to allow increased aeration). All cultures were grown at 30 ℃ for ~8 hours or until $OD_{600nm}$ = 0.6 was reached. Subsequently, controls and the library were induced by addition of IPTG. Cultures were grown for an additional 4 hours or until $OD_{600nm}$ = ~1.

To prepare cells for the selection condition, 1 mL of each culture was pelleted in 2 mL microcentrifuge tubes and washed three times with M9 Wash Buffer. After wash, cells were finally re-suspended in 1 mL M9 Wash Buffer. Cells were diluted with M9 Wash Buffer to a final cell concentration of ~$10^7$ cells/mL. Ten 100 μL aliquots of this cell suspension were plated on separate M9 Selection Plates supplemented with 2 g/L cyclohexanone and incubated at 30 ℃ for 48 hours. The colony growth was monitored periodically. Selected colonies were re-streaked onto fresh selection media and again incubated to obtain single colonies. Colonies were favored based on initial time of observable growth and size. From selection, 26 single colonies were cultured in liquid media to extract plasmids using QIAprep Spin Miniprep kit (Qiagen). Sequencing results highlighted variant on plasmid pLS210 (DTNP) which appeared four times in contrast to majority of variants which appeared once and no more than twice. To re-confirm growth phenotype conferred, variant plasmids were re-transformed individually into MX304 and subjected to identical selection protocol. Due to the superior growth conferred by DTNP compared to other variants and preliminary activity assays, only variant encoding the CHMO DTNP was extensively characterized.

**Expression and Purification of CHMO Wild-type and Variants.** Enzyme expression of pQElac based plasmids used *E. coli* BL21 (DE3) containing the plasmids encoding for the CHMO wild-type or variants. Enzyme expression of pRSF based plasmids used *E. coli* strain MX203. Expression was carried out by inoculation of 50 mL 2xYT media supplied with 100 μg/mL ampicillin and an overnight culture. Cells were grown at 37 ℃ in baffled shaking flasks and were induced at an $OD_{600nm}$ of 0.6–0.8 with 0.25 mM IPTG. The 50 mL main culture was then incubated at 25 ℃ and 250 rpm for 24 h. The expression was stopped by harvesting the cells (centrifugation at 4 ℃, 4000 g for 15 min). Cell pellet was resuspended in HisPur™ Ni-NTA equilibration euffer (Thermo Fisher Scientific) and mixed with 1 mL of 0.1 mm glass beads (Biospec) and homogenized using a benchtop homogenizer (FastPrep-24, MP Biomedicals). Cell debris was separated from the crude extract by centrifugation at 4 ℃, 20,000 g for 15 min. Protein purification was performed using HisPur™ Ni-NTA Protein Miniprep (Thermo Fisher Scientific) according to the manufacturer's instructions. The histidine-tagged protein was eluted in HisPur™ elution buffer (50 mM sodium phosphate buffer pH 7.5, 300 mM NaCl, 300 mM imidazole). The concentrations of purified protein were quantified by Bradford assay using BSA as standards. For stock protein, 20% (v/v) glycerol was added; and the protein was stored at −80 ℃ for future use or used immediately for temperature gradient incubation specific activity assay.

**Temperature Gradient Incubation Specific Activity Assay.** To determine thermostability, residual specificity activity values were determined for CHMO variants according to previously described methods[7]. Purified and diluted CHMO variants (~0.03-0.05 mg/mL) were incubated in sodium phosphate buffer (50 mM, pH 7.7) at various temperatures (25 ℃, 30 ℃, 40 ℃, and 45 ℃) for 10 min within PCR tubes using a gradient temperature controlled Thermocycler (Biorad). Afterwards the samples were immediately placed on ice for a 5 minute incubation period and activity was subsequently measured at 25 ℃ using NADPH assay (0.1mM NADPH, 5 mM cyclohexanone (1 M stock was dissolved in ethanol), 50 mM sodium phosphate buffer pH 7.7, 3-5 μg/mL protein).

**Steady State Kinetic Analyses for Cyclohexanone and NAD(P)H.** Catalytic efficiency ($k_{cat}/K_m$) for NAD(P)H (Table 2 and Table S3) were performed according to previously described methods[8]. The reaction mixture containing 50 mM Tris-Cl buffer (pH 9.0), 1 mM cyclohexanone and varied NAD(P)H concentrations from 0.02 mM to 4 mM was incubated at 25 ℃. For kinetic characterization in Table S2, concentrations of substrates and cofactors were changed (0.002-0.1 mM for cyclohexanone; 0.005-0.2 mM for NADPH) to allow the determination of the different steady-state kinetic parameters, the reaction was performed in 50 mM sodium phosphate (pH 7.7) at 25 ℃. All the reactions were initiated by addition of an appropriate amount of the enzyme (purified protein from −80 °C stock) and the kinetic parameters were measured by monitoring the NAD(P)H consumption at 340 nm in 96-well plate.

***Ac* CHMO Homology Modeling.** The model of WT *Ac* CHMO with cofactors FAD and NADPH bound was generated with Rosetta CM[9,10]. Threading templates with high sequence identity and co-crystallized cofactors were identified through BLASTP search of the Protein Data Bank with *Ac* CHMO as the query[11,12]. Crystal structures of CHMO from *Rhodococcus sp.* (PDB: 4RG3, 3GWD, 3GWF, 3UCL), which shares 57.8% sequence identity to *Ac* CHMO, were selected as input models[14–16].The Rosetta CM protocol consisted of repeated rounds where: the target *Ac* CHMO sequence was threaded onto the template structure based on MAFFT sequence alignment, segments of the protein structure were constructed through insertion of fragments drawn from the library provided by the templates through Monte Carlo evaluation, followed by minimization to relax the final output[17]. 1,500 homology modeling trajectories were completed, the output structure with the most favorable total Rosetta energy was selected as the representative model for all further analysis. Point mutations for the *Ac* CHMO variants were produced through 1,000 further Rosetta docking simulations on the homology model with backbone flexibility, side chain repacking, and ligand minimizations; final models were selected by lowest total Rosetta energies. The NADH binding poses were prepared by deleting atoms of the ribose 2’ phosphate group on the existing NADPH models prior to the Rosetta Design trials.

The *Ac* CHMO model shows that NADPH binds in an extended conformation with the nicotinamide ring tucked into a small binding pocket against the FAD flavin. The NADPH binding mode is characterized by the conserved Rossman fold with β-α-β secondary structure motifs enclosing the cofactor. The primary interactions predicted to hold the NADPH in place include: Q190 side-chain amide which contacts the NADPH carboxamide oxygen, W490 indole which forms a hydrogen bond to the nicotinamide ribose hydroxyl, backbone polar interactions at the N-terminus of the Rossman α-helix and hydrogen bonding from the T189 hydroxyl to the pyrophosphate, a salt-bridge from K326 extending from a loop across the substrate channel to the pyrophosphate, and R207 guanidinium forming a salt-bridge to the ribose 2’ phosphate group and packing against the adenosine ring. The side of the NADPH facing away from the Rossman fold is marginally exposed, with the control loop defined as residues 489-505 lightly packing against the cofactor. Since NADH differs from NADPH only in the absence of the 2’ phosphate group and is capable of establishing the same set of binding interactions along the pyrophosphate and nicotinamide ring, we postulate that *Ac* CHMO’s strict specificity for NADPH is driven by the R207 contact spanning from the end of the second Rossman β-strand stabilizing the adenosine tail of the cofactor for optimal packing dynamics with the control loop. The FAD is held opposite of the nicotinamide cofactor with the flavin group facing the nicotinamide ring and is tightly bound through polar contacts throughout the adenosine tail, ribose, pyrophosphate, ribitol, and flavin carbonyls. The numerous hydrogen bonds restrict FAD mobility, position the flavin for efficient hydride transfer with the nicotinamide cofactor or substrate, and prevent the FAD release. The cyclohexanone binding pocket is adjacent to the nicotinamide ring, it is proposed that some degree of protein and NADPH flexibility is required to allow the substrate to move close enough to the FAD for electron transfer

**Molecular Dynamics Simulations.** MD simulations of the *Ac* CHMO variants were completed with PMEMD from the AMBER 18 package utilizing the ff14sb force field and 8 Å Particle Mesh Ewald real space cutoff[18–21]. Cofactor parameters were obtained from the AMBER parameter database and protonation states of titratable residues were determined with the H++ webserver[22–24]. The TLEAP program was utilized to solvate the complexes with TIP3P water molecules in a truncated octahedron with 10 Å buffer and neutralizing Na+/Cl- counter-ions. The *Ac* CHMO systems were minimized in two stages, first with 2,500 steps of steepest decent and 2,500 steps of conjugate gradient where all non-hydrogen solute atoms were restrained with a 20 kcal $mol^{-1}$ $Å^{-2}$ force to relieve solvent clash. The second stage minimization to remove solute steric clashes was run with the same cycle settings and restraints removed. Heating from 0 K to 300 K was performed over 0.5 ns with 10 kcal $mol^{-1}$ $Å^{-2}$ restraints on all non-hydrogen solute atoms under NPT conditions at 1 atm pressure with Langevin thermostat and 1 fs timestep. Solvent density equilibration over 5 ns with 5 kcal $mol^{-1}$ $Å^{-2}$ restraints on all solute atoms and an unrestrained 10 ns equilibration using 2 fs timestep to clear remaining structural artifacts followed the heating stage. Production MD trajectories were each carried out for 400 ns with 2 fs timestep, SHAKE restraints on hydrogens, NVT ensemble, Langevin thermostat with collision frequency 1.0 $ps^{-1}$, and periodic boundary conditions.

**Thermostability and Cofactor Binding Analysis.** Metastable conformations sampled by the *Ac* CHMO systems and representative frames for the discovered states were identified through PCA dimensionality reduction and K-Means clustering[25,26]. To understand the mechanisms behind the enhanced thermostability of GV and WT *Ac* CHMO, we compared MD trajectories of GV and WT *Ac* CHMO featurized on minimum heavy atom distance between the residues at 245 or 288 to neighboring residues having any atom within 5 Å contact. The residue positions around 245 are selected as: 62, 239, 242, 243, 244, 245, 246, 247, 250, 433, 434, 437, and 510. The residue positions around 288 are defined as: 280, 281, 286, 287, 288, 289, 290, 294, 482, 483, and 484. The distance array was standardized to zero mean and unit variance and transformed to lower dimensional space with components maintaining maximal variance through PCA. K-means clustering was performed to discretize the sampled conformations projected onto the free energy landscape of the first two PCA components into metastable states. The optimal number of clusters was selected by the elbow heuristic where clustering over a range of K values, from one to nine here, is completed and the sum of squared distances from the sample points to their assigned cluster center is computed. The value of K where the sum of squared distances decrease becomes linear is selected as optimal and indicates that increasing K further will result in over-fitting. Thermostability was evaluated through the metric of Rosetta residue energies, a score quantifying the favorability of van der Waals, electrostatic, and geometric interactions that functions as a surrogate for predicted stability, summed over all residues within 5 Å of the mutated positions[27]. 200 frames were extracted from the most populated metastable state for each sample and scored with Rosetta, the conformation with minimum distance to the cluster center was chosen for illustration.

Cofactor binding comparisons were completed between WT *Ac* CHMO, DTN CHMO, and DTNP CHMO with either NADH or NADPH bound. Protein backbone flexibility over the trajectories was measured through alpha-carbon RMSF. Hydride transfer potential was recorded as the distance between the nicotinamide C4 to FAD N5. Metastable FAD binding conformations were established through featurization on minimum heavy atom distance to residues with any atom within 4 Å contact and similar PCA/K-means analysis as described above. The residue positions neighboring FAD include: 12, 13, 15, 16, 17, 36, 37, 44, 45, 46, 48, 49, 51, 56, 57, 58, 63, 109, 110, 140, 141, 142, 390, 426, 434, 435, 436, and 439. MDTraj and cpptraj were utilized for trajectory analysis, data processing was completed with the NumPy and scikit-learn packages[28–31]. PyMol was used to illustrate the structures (Schrödinger LLC, 2020).

**Bioinformatic Analysis.** Homologous sequences to WT *Ac* CHMO were identified through BLASTP search over the non-redundant protein sequence database[13]. Sequences were filtered on the criteria of query coverage over 70% and sequence identity between 40% to 97%, resulting in 998 hits. MSA of the matched sequences to the template *Ac* CHMO was completed with MAFFT[17]. Residue frequencies for each of the 20 canonical amino acids were computed at every position in the multiple sequence alignment.

**B. Supplemental Tables and Figures**

**Table S1. Strains and Plasmids used in this study**

| **Strains** | **Description** | **Reference** |
|---|---|---|
| XL-1 blue | Cloning strain | Stratagene |
| BL21 (DE3) | Protein expression strain | Invitrogen |
| BW25113 | *E. coli* F-, DE(araD-araB)567, lacZ4787(del)::rrnB-3, LAM-, rph-1, DE(rhaD-rhaB)568, hsdR514 | Datsenko *et al.*[4] |
| DH10β | Electrotransformation strain | Invitrogen |
| JW2279-3 | BW25113 Δ*nuoF::kan* | Coli Genetic Stock Center |
| JW1095-1 | BW25113 Δ*ndh::kan* | Coli Genetic Stock Center |
| JW5713-1 | BW25113 Δ*ubiC::kan* | Coli Genetic Stock Center |
| JW1594-1 | BW25113 Δ*pntB::kan* | Coli Genetic Stock Center |
| JCL166 | BW25113/F′ [*traD36 proAB*$^+$*lacI*$^q$ZΔM15 (Tet$^r$)] Δ*ldhA* Δ*adhE* Δ*frdBC* | Atsumi *et al.*[32] |
| MX203 | BW25113 Δ*pgi* Δ*edd* Δ*qor* Δ*udhA::kan* | Maxel *et al.*[2] |
| MX301 | JCL166 Δ*nuoF* Δ*ndh::kan* | This study |
| MX302 | JCL166 Δ*nuoF* Δ*ndh::kan* + pLS101 | This study |
| MX303 | JCL166 Δ*nuoF* Δ*ndh* Δ*ubiC::kan* + pLS101 | This study |
| MX304 | JCL166 Δ*nuoF* Δ*ndh* Δ*ubiC* Δ*pntB::kan* + pLS101 | This study |
| **Plasmids** | **Description** | **Reference** |
| pCP20 | Temperature-inducible yeast Flp recombinase gene controlled by λ*c*I*ts857* in a temperature-sensitive replicon, Amp$^r$ | Datsenko *et al.*[4] |
| pQElac | Amp$^r$ ; ColE1 ori; $P_{\mathrm{LlacO1}}$. Expression vector | Li *et al.*[33] |
| pLS101 | pRSF $P_{\mathrm{BAD}}$::*Lb nox*, Spec$^r$ | Maxel *et al.*[2] |
| pLS102 | pRSF $P_{\mathrm{BAD}}$::*TP nox* (*Lb nox* G159A-D177A-A178R-M179S-P184R), Spec$^r$ | Maxel *et al.*[2] |
| pLS201 | pRSF $P_{\mathrm{BAD}}$:: *Ac chnB*, Spec$^r$ | This study |
| pLS202 | pRSF $P_{\mathrm{BAD}}$:: *Ac chnB* T415C, Spec$^r$ | This study |
| pLS203 | pRSF $P_{\mathrm{BAD}}$:: *Ac chnB* Error Prone-PCR library, Spec$^r$ | This study |
| pLS204 | pRSF $P_{\mathrm{BAD}}$:: *Ac chnB* A245G-A288V, Spec$^r$ | This study |
| PLS205 | pRSF $P_{\mathrm{BAD}}$:: *Ac chnB* A245G-A288V-T415C, Spec$^r$ | This study |
| pLS206 | pQElac 6xHis *Lb nox*, Amp$^r$ | This study |
| pLS207 | pQElac 6xHis *TP nox* (*Lb nox* G159A-D177A-A178R-M179S-P184R), Amp$^r$ | This study |
| pLS208 | pQElac 6xHis *Ac chnB*, Amp$^r$ | This study |
| pLS209 | pQElac 6xHis *Ac chnB* S208-K326-K349-T378 NNK library, Amp$^r$ | This study |
| pLS210 | pQElac 6xHis *Ac chnB* L143P-S208D-K326T-K349N, Amp$^r$ | This study |
| pLS211 | pQElac 6xHis *Ac chnB* S208D-K326T-K349N, Amp$^r$ | This study |
| pLS212 | pQElac 6xHis *Ac chnB* S186P-S208E-K326H-K349R, Amp$^r$ | This study |

Abbreviations indicate source of genes: *Lb*, *Lactobacillus brevis*, *Ac Acinetobacter sp. NCIMB 9871*

**Table S2. Kinetic characterization of CHMO and GV.**

| Enzymes | NADPH | | | Cyclohexanone | | |
|---|---|---|---|---|---|---|
| | $K_m$( μM) | $k_{cat}(s^{-1})$ | $k_{cat}/K_m$ ( $\mu M^{-1}s^{-1}$) | $K_m$( μM) | $k_{cat}(s^{-1})$ | $k_{cat}/K_m$ ( $\mu M^{-1}s^{-1}$) |
| Wild type | 36.6 ± 8.36 | 13.5 ± 0.55 | 0.37 | 4.09 ± 1.81 | 12.9 ±0 .86 | 3.15 |
| GV | 43.3 ± 20.9 | 9.67 ± 1.23 | 0.22 | 5.64 ± 0.48 | 11.6 ±1.55 | 2.05 |

Activities were measured as described in Methods. Briefly, kinetic constants determined for NADPH and cyclohexanone are performed at pH 7.7 and 25 ℃ for both wild-type CHMO and GV. Kinetic parameters are the average of three or two independent experiments with numbers after ± stand for one standard deviations for NADPH or cyclohexanone, respectively.

**Table S3. Kinetic characterization of CHMO PEHR.**

| CHMO Variants | $k_{cat}/K_m$ ($mM^{-1}s^{-1}$) | |
|---|---|---|
| | NADH | NADPH |
| Wild type | 20.8 ± 3.51 | 400 ± 25.8 |
| PEHR | 21.5 ± 1.69 | 21 ± 4.67 |

The kinetic characterizations for cofactors were measured in 50 mM Tris-Cl buffer (pH 9.0) with 1 mM cyclohexanone and varied NAD(P)H concentrations from 0.02 mM to 4 mM at 25 ℃. All the kinetic parameters are the average of three independent experiments with numbers after ± stand for one standard deviations. CHMO PEHR (S186P-S208E-K326H-K349R) is a previously reported NADH active CHMO variant[8]. This variant was constructed through the addition of several individually beneficial mutations discovered in rational design. Individual characterization of $k_{cat}$ and $K_M$ was not achieved because the enzymes could not be saturated in the range of cofactor concentrations tested.

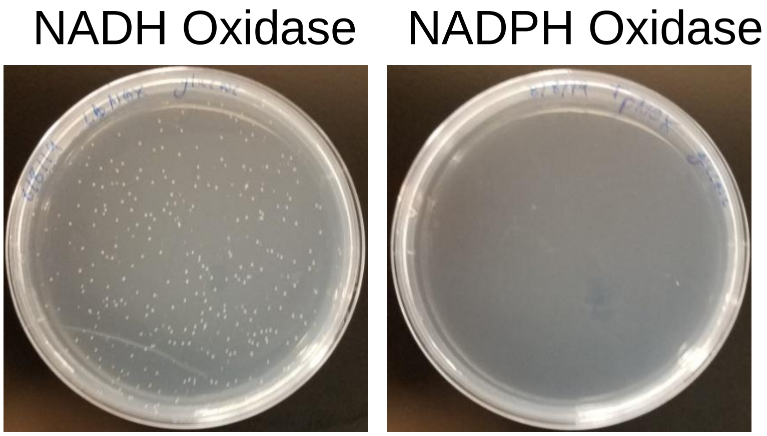


**Figure S1.** MX304 growth behavior with oxidase complementation on solid minimal media with glucose as sole carbon source. Growth restoration was observed when expressing NADH-dependent oxidase (Left). Little to no growth observed with NADPH-dependent oxidase expression (Right).

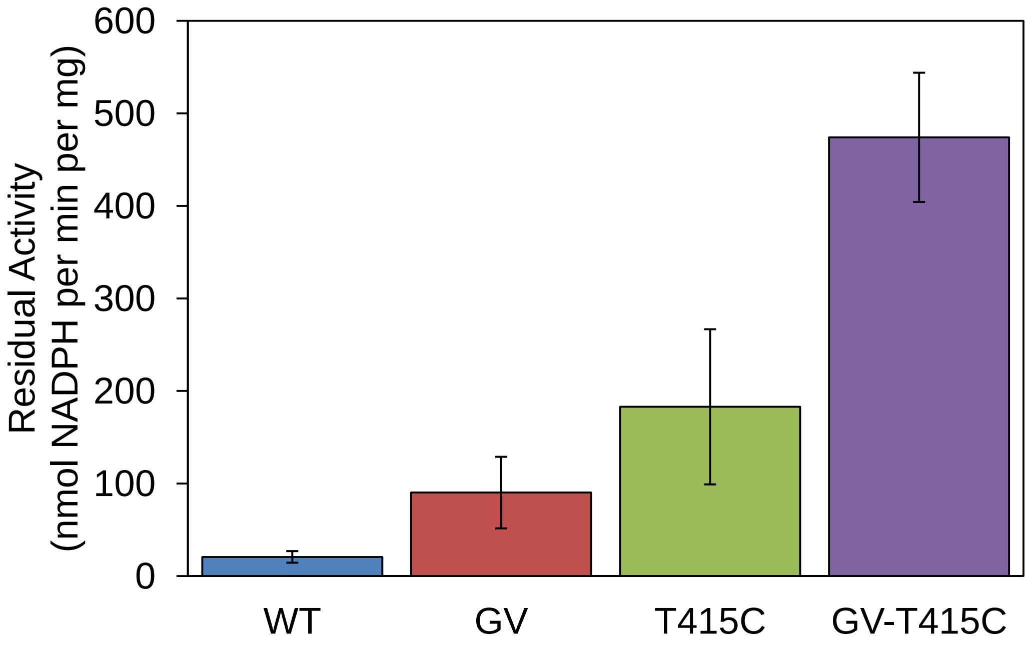


**Figure S2.** Residual specific activity of CHMO variants after ten-minute incubation at 45 ℃. Variants GV and T415C displayed 4.4 and 8.8-fold improved residual activity over WT. Variant combining the stabilizing mutations GV and T415C significantly increased residual activity 23-fold suggesting a synergistic effect on protein stability.

After selection at 42 ℃, the combination of A245G and A288V was found to confer substantial improvements in residual activity at 45 ℃ (Figure S2). We sought to explore potential synergy between the mutations in variant GV and T415C, a single-point mutant described by Schmidt et al. with significant improvements in long-term stability[7]. Introduction of T415C, a free cysteine, in addition to selected mutations, yields a triple mutant with greatly improved activity and stability at 45 ℃ relative to variant GV or T415C alone. The additive effect observed for the three distal mutations is likely a consequence of their spatial isolation, where changes independently address local instabilities that could result in protein unfolding[34]. Although structure guided engineering of thermostability is encumbered by the need to comprehensively account for interactions between the entirety of the protein and solvent, synergistic random mutations can be more frequently encountered because localized perturbations can be distributed throughout the structure and in the absence of direct interaction.

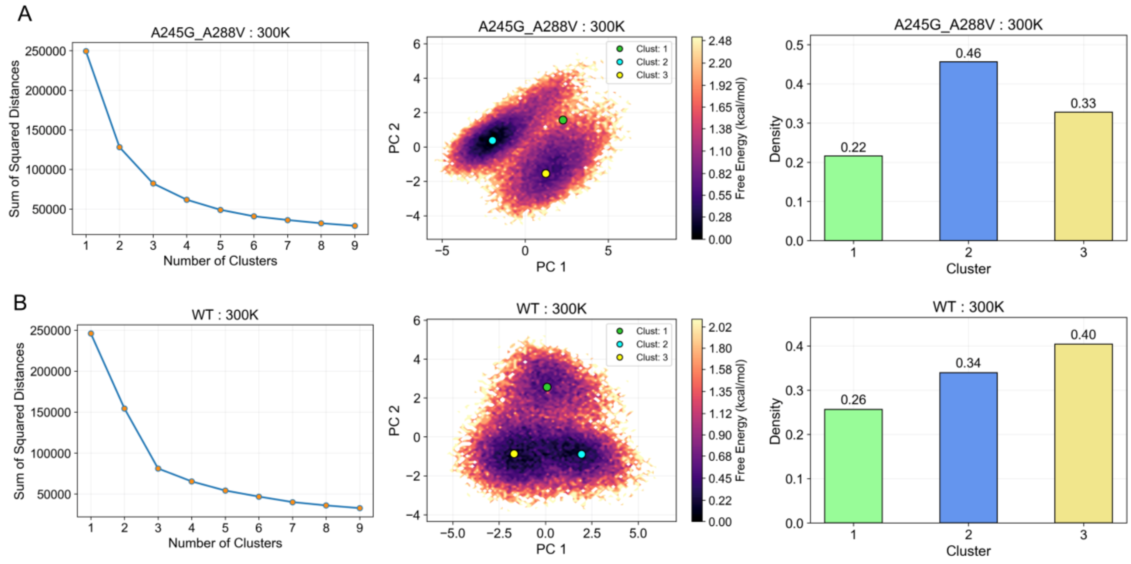


**Figure S3.** Free energy landscapes for GV and WT CHMO. **A)** K-means clustering elbow heuristic to determine the optimal number of clusters, free energy landscape projected on first 2 principal components and cluster centers, cluster populations. **B)** WT CHMO K-means clustering, free energy landscape, and cluster populations.

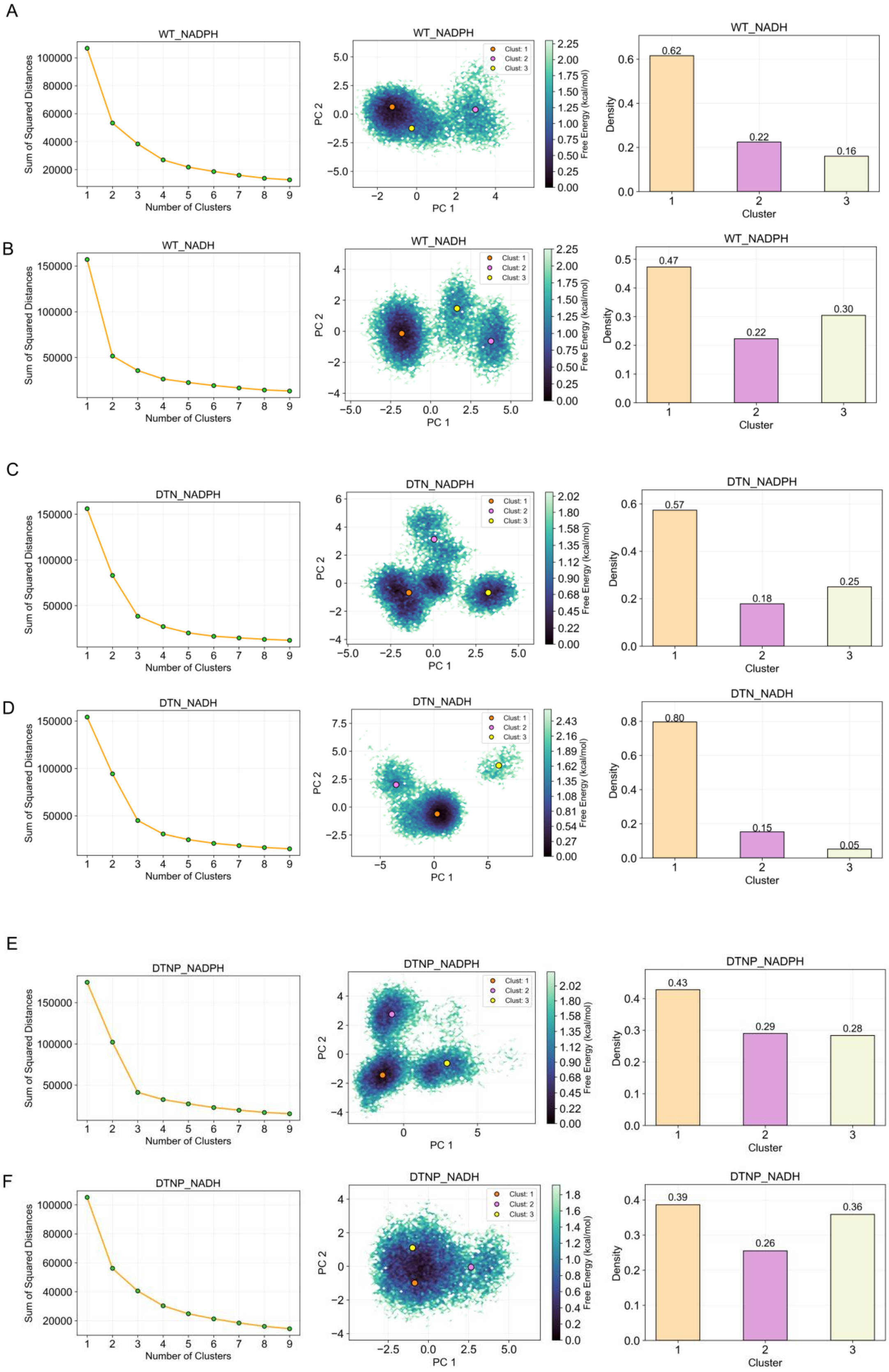
A
WT_NADPH
Sum of Squared Distances
Number of Clusters
WT_NADPH
Clust: 1
Clust: 2
Clust: 3
PC 1
PC 2
Free Energy (kcal/mol)
WT_NADH
Density
Cluster
0.62
0.22
0.16
B
WT_NADH
Sum of Squared Distances
Number of Clusters
WT_NADH
PC 1
PC 2
Free Energy (kcal/mol)
WT_NADPH
Density
Cluster
0.47
0.22
0.30
C
DTN_NADPH
Sum of Squared Distances
Number of Clusters
DTN_NADPH
PC 1
PC 2
Free Energy (kcal/mol)
DTN_NADPH
Density
Cluster
0.57
0.18
0.25
D
DTN_NADH
Sum of Squared Distances
Number of Clusters
DTN_NADH
PC 1
PC 2
Free Energy (kcal/mol)
DTN_NADH
Density
Cluster
0.80
0.15
0.05
E
DTNP_NADPH
Sum of Squared Distances
Number of Clusters
DTNP_NADPH
PC 1
PC 2
Free Energy (kcal/mol)
DTNP_NADPH
Density
Cluster
0.43
0.29
0.28
F
DTNP_NADH
Sum of Squared Distances
Number of Clusters
DTNP_NADH
PC 1
PC 2
Free Energy (kcal/mol)
DTNP_NADH
Density
Cluster
0.39
0.26
0.36

**Figure S4.** Free energy landscapes for DTN, DTNP, and WT CHMO with NADH or NADPH bound. **A)** WT with NADPH bound **B)** WT with NADH bound **C)** DTN with NADPH bound **D)** DTN with NADH bound **E)** DTNP with NADPH bound **F)** DTNP with NADH bound
